# Chemical degumming increases larvae size and facilitates the commercial production of Lumpfish (*Cyclopterus lumpus*) eggs

**DOI:** 10.1101/583641

**Authors:** Craig Pooley, Mia Berwick, Carlos Garcia de Leaniz

## Abstract

Many fishes produce adhesive eggs that confer protection from currents and predators in the wild, but that are more difficult to disinfect and aerate under aquaculture conditions. Removing egg adhesiveness (‘degumming’) has proved beneficial in the culture of many fish, and a recent gap analysis identified this as a potential way of increasing hatching success and minimize the risk of infectious diseases in the culture of lumpfish (*Cyclopteurs lumpus*), a novel species to aquaculture. We tested the efficacy of the enzyme alcalase (0.02%, 0.2%, 2%) as a degumming agent for lumpfish eggs, and examined its effects on hatching success, survival, and larvae size under laboratory and commercial conditions. A five-minute exposure to 0.2% and 2% alcalase decreased chorion thickness by 14% and resulted in 61-75% degumming rates, without any negative effects on hatching rate, larval survival, or incidence of embryo malformations. Degummed eggs hatched earlier than controls and resulted in larger larvae, which may confer some benefits under aquaculture conditions. A cost-benefit analysis indicates that the benefits of egg degumming compensate the costs of chemical treatment under most conditions, and that the optimal alcalase concentration is around 0.2%. We therefore recommend egg degumming as a way of making the lumpfish industry more efficient and sustainable.

**Statement of relevance:** Improving the commercial production of lumpfish

## 1. Introduction

Many fish species lay eggs that become adhesive in contact with the water, which is thought to be adaptive as it increases embryo survival (Rizzo et al., 2002). For example, some freshwater fish produce sticky eggs that cling to the substrate to avoid being washed away with the current, while other species may produce adhesive eggs to form discrete clumps that are easier to tend, or that stick to the vegetation to decrease predation risk or increase oxygen supply (Riehl & Patzner,1998). However, the same adhesiveness that benefits eggs in the wild may hamper their viability under aquaculture conditions, where eggs are typically reared at very high densities and, if untreated, are prone to suffer from fungal infections and other diseases (Ringle et al, 1992).

The benefits of removing the adhesiveness of the eggs (‘degumming’) in fish farming have long been known (Rottman et al., 1991; Billard et al. 1995). These include improved aeration, reduced risk of fungal diseases, more effective disinfection, and less effort associated with egg count and husbandry (Ringle et al, 1992). Egg degumming has proved beneficial for the culture of many freshwater fishes, including cyprinids (Linhart et al, 2003; El-Gamal & El-Greisy, 2008; Žibienė et al., 2017), tench (*Tinca tinca*, Linhart et al., 2000; Gela et al., 2003; Kujawa et al., 2010), walleye (*Sander vitreus*, Krise et al., 1986), pikeperch (*Sander lucioperca,* Demska-Zakes et al., 2005), barbels (Al Hazzaa & Hussein, 2003), sturgeons (Pšenička, 2016), and catfishes (Isaac & Fries, 1991; Ringle et al., 1992; Linhart et al., 2003; Asraf et al., 2013; Muchlisin et al., 2014; Kareem, et al., 2017). Although less common in marine aquaculture, egg degumming has also proved effective for anadromous and marine species (white sturgeon, *Acipenser transmontanus*; Kowtal et al, 1986; ballan wrasse, *Labrus bergylta;* Grant et al., 2016).

Naturally sticky fish eggs have been chemically treated with many different degumming agents with various degrees of success, including urea (Kowtal et al., 1986), milk (Billard et al., 1995; Linhart et al., 2000), pineapple juice (Thai & Ngo, 2004), clay (Siddique et al., 2016), talc (Linhart et al. 2002), tannic acid (Demska-Zakes et al., 2005), sodium chloride (Kowtal et al., 1986) and sodium sulphite (Isaac & Fries, 1991) among others. Glycoproteins and glycosaminoglycans present in the egg chorion are the molecules responsible for making fish eggs sticky (Riehl & Patzner,1998), and for this reason enzymes that can break down peptide bonds, such as papain (Isaac & Fries, 1991), trypsin (Linhart et al., 2002) and various endo-proteases (Linhart et al., 2003; El-Gamal & El-Greisy, 2008), have proved particularly effective as degumming agents. Among these, alcalase (a commercial endo-protease with broad specificity obtained from the bacterium *Bacillus licheniformis)* has been effective in both freshwater (Linhart et al., 2003) and marine species (Grant et al., 2016). However, egg degumming also poses risks and could easily damage the fragile fish embryo (Pšenička, 2016). Given that species differ markedly in the structure and chemical composition of the egg chorion (Riehl & Patzner, 1998), no single degumming method is likely to be suitable in all cases.

One novel species to Aquaculture that has adhesive eggs and whose incubation in hatcheries could benefit from egg degumming is the lumpfish, *Cyclopterus lumpus* (Powell et al., 2018a). Large scale commercial production of lumpfish only started recently but has increased exponentially over the last few years as the species is increasingly being used as cleaner fish to control parasitic sea lice in salmon farming (Powell et al., 2018a). Different lumpfish populations have different fecundities and size at maturation (Whittaker et al., 2018), but an analysis identified egg degumming as a potential way of increasing hatching success and minimizing the risk of infectious diseases during embryo development (Powell et al., 2018a). Under natural conditions, lumpfish eggs become sticky following exposure to divalent calcium and magnesium in seawater (Lönning et al., 1984). Following fertilization, males create funnel-like depressions in the egg mass and puff water and vigorously fan the eggs to facilitate gaseous exchange and the removal of nitrogenous waste, such paternal care being maintained throughout the incubation period (Powell et al., 2018b). In contrast, lumpfish eggs in hatcheries may be left to incubate in large clumps or shaped into a thin monolayer and reared in upwelling incubators (Powell et al., 2018a). Hatching success is very variable in culture and it is not uncommon to lose entire egg batches due to poor oxygenation and subsequent spread of bacterial and fungal diseases (Powell et al., 2018a). Since a female can typically produce in excess of 100,000 eggs (Davenport, 1985; Powell et al., 2018b), even a modest increase in hatching rate would yield a significant improvement in the number of larvae produced.

As with ballan wrasse (Grant et al., 2016), a small-scale pilot trial showed recently that alcalase could also be effective at degumming lumpfish eggs (Powell et al., 2018a), but the side effects and most efficient concentrations were not investigated. Thus, the aim of this study was to test the potential benefits of alcalase treatment to remove the adhesive layer of the lumpfish eggs and examine any potential impacts on embryo survival, timing of hatching, growth, and incidence of malformations. To this end, two experiments were conducted, one under controlled laboratory conditions, where accurate counts could be made of daily mortalities in replicated mini-incubators, and one under commercially relevant conditions in order to validate the results obtained in the laboratory.

## 2. Materials and Methods

### 2.1. Source of eggs and artificial fertilization

Wild lumpfish were obtained from commercial fishermen operating in Dorset (50° 44’ N; 2° 20’ W) and Guernsey (49° 27’ N; 2° 31’ W) during 2016 and 2017 and were transferred to a quarantine unit at the Centre for Sustainable Aquatic Research (CSAR, Swansea University) where they were kept in 1.5 m3 circular tanks at 32 ppt salinity, 13°C ± 0.3°C, and 8:16 L:D photoperiod. Broodstock were fed a mixture of frozen blue mussels, squid, krill and whitebait.

Eggs were stripped from anaesthetized females (2-phenoxyethanol, concentration 0.03 ml/litre) and milt was collected from euthanized males as described in Powell et al. (2018a). Sperm motility was assessed by activation of a small sample with seawater, followed by observation under the microscope at x40 magnification. Sperm was diluted with a milt extender (AquaBoost SpermCoat, Cryogenetics AS, Norway) at a 1:1 ratio and maintained refrigerated at 4°C for up to 14 days. Artificial fertilization was carried out by adding 2-3 ml of extended milt to a batch of eggs (~600 g), followed by gentle agitation for 1 min, activation with ozone-treated seawater, further agitation for 2 min, and rinsing with clean seawater. Egg hardening was completed after c. 30 min at 10°C.

### 2.2. Laboratory (in vitro) degumming trial

Three liquid alcalase (*Bacillus licheniformis*; 126741; VWR, UK) solutions were prepared using 0.2 μm filtered seawater as a diluent to achieve concentrations of 0.02%, 0.2% and 2.0% along with a control treatment (0% alcalase). These were maintained in the dark at 4°C for a maximum of 48h prior to use. Immediately following sperm activation, and prior to egg hardening, 12 batches of fertilized eggs (n = 104 ± 10 eggs) from a single female were exposed for 5 min to one of the four alcalase concentrations, gently inverting the vial to ensure uniform exposure for all eggs. Following alcalase exposure, eggs were allocated at random to 12 × 0.5 L upwelling vertical incubators constructed from 750 mL plastic funnels fitted with a 500 μm mesh “cradle” to hold the eggs. Flow was maintained at 1 Lmin-1 (± 0.05 L), and water temperature was maintained at 8.0 °C ± 0.1 °C during the 37-day duration of the trial. Eggs were disinfected with 30 ppm Pyceze (Novartis Pharmaceuticals UK Ltd) for 20 minutes every 5 days.

Eggs were inspected 48 h post-fertilisation (morula stage) to assess their viability and estimate fertilization success (Powell et al., 2018b). Degumming efficiency was assessed by counting the number of single eggs (i.e. those not in a clump, bound together by their adhesive chorion) and dividing by the total number of eggs in the sample. They were then photographed using a Leica DFC290 digital camera (Leica Microsystems Ltd, UK) mounted to a Nikon SMZ 800 Zoom Stereomicroscope at X20 magnification. Mean chorion thickness (±10 μm) was obtained from photomicrographs of six eggs per replicate (n=18 per treatment) at 2-4 equidistant points of the egg, as described in Songe et al. (2016).

Hatched larvae were collected daily from individual incubators and preserved in 10% formalin (HT501128; Sigma-Aldrich, UK) to screen for embryo malformations. Hatching success was calculated by dividing the number of hatched larvae by the number of viable eggs (i.e. excluding non-viable eggs that failed to develop after 24 hrs). A sample of fixed larvae (n=6 per replicate, n=18 per treatment) were photographed and the standard length was calculated from the tip of the snout to the caudal peduncle (resolution ± 10μm). The incidence of embryo deformities such as spinal scoliosis (Gapasin et al., 1998; Merchie et al., 1997), and operculum and sucker deformities was recorded.

### 2.3. Degumming under commercial conditions

A 2% alcalase concentration was chosen for the large commercial trial, as this had shown promising results in a small-scale pilot study (Powell et al., 2018a). A solution was prepared by adding 10 ml of alcalase to a 20-litre beaker containing 490 ml of filtered sea water. Fertilised lumpfish eggs (590 ± 10 g) were added to the beaker and aerated. For the control treatment, the same amount of eggs was added to 500 ml of filtered sea water. After 5 minutes, eggs were rinsed with 1 litre of filtered sea water and transferred to 70 litre hopper incubators with a 20 L/min upwelling flow and a 500 µm screen mesh to keep the eggs off the bottom. Salinity was maintained at 32 ppt and water temperature at 9.0 ± 2.0 °C. Eggs were treated with Pyceze every five days, as above. Approximately 30 eggs were sampled from each clump using a grid system to ascertain viability and degumming efficiency, as described above. During the peak of mass hatching, 100 larvae per batch were reared in triplicate in 30 litre tanks and monitored for eight weeks under the same commercial conditions (temperature 9.0 ± 2.0 °C) and fed enriched *Artemia* and dry diet. Mortalities were recorded daily for each replicate.

### 2.4. Statistical analysis

We used R v3.4.1 (R Core Team, 2017) for statistical analysis. Degumming efficiency, survival, and incidence of embryo malformations (all proportion data) were analysed via Generalized Linear Mixed-Effect Modelling (GLMM) using the R package “*lme4*” (Bates et al., 2015) and a binomial error distribution, with alcalase concentration as a fixed factor and incubator ID as a random effect. Hatching success was modelled as a function of time and alcalase concentration as predictors, and incubator ID as a random effect, using the packages “*survival*” (Therneau, 2015) and “*FrailtyPack*” (Rondeau et al., 2012) to account for unmeasured heterogeneity. The effect of alcalase concentration on chorion thickness and larvae size at hatching was tested by Linear Mixed Modelling (LMM) using alcalase concentration as a fixed effect, and incubator identity as a random effect using the *lme4 and lmerTest* (Kuznetsova et al., 2017) R packages. We compared models using untransformed and log10 transformed alcalase concentrations, and also treating alcalase concentration as a continuous or discrete predictor. Inspection of AIC values indicated that treating alcalase concentration as a factor resulted in the most plausible models.

### 2.5. Ethics Statement

This study adhered to the ARRIVE guidelines and was approved by Swansea University Animal Welfare Research Body (permit IP-1516-16)

## 3. Results

### 3.1. Efficacy of alcalase as a degumming agent

Alcalase was effective at separating (degumming) lumpfish eggs at concentrations of 0.2% and 2% for 5 minutes, but not at 0.02% (Figure 1, glmer: estimate = 1.97, SE = 0.226, Z value = 8.724, *P* < 0.001). Compared to controls, eggs exposed to alcalase concentrations of 0.2 and 2% displayed mean degumming rates of 61% (±12) and 75% (±21), respectively. The degumming effect of alcalase was confirmed by measurements of chorion thickness, which decreased significantly with increasing alcalase concentration (Figure 2; lmer; *F_3,11.852_* = 11.102, *P* < 0.001). Thus, compared to controls (mean chorion thickness = 68.59 ± 2.91µm), chorion thickness decreased by 8.77 μm (±1.91) at 0.2% and by 10.24 μm (±1.93) at 2% alcalase.

**Figure 1.**
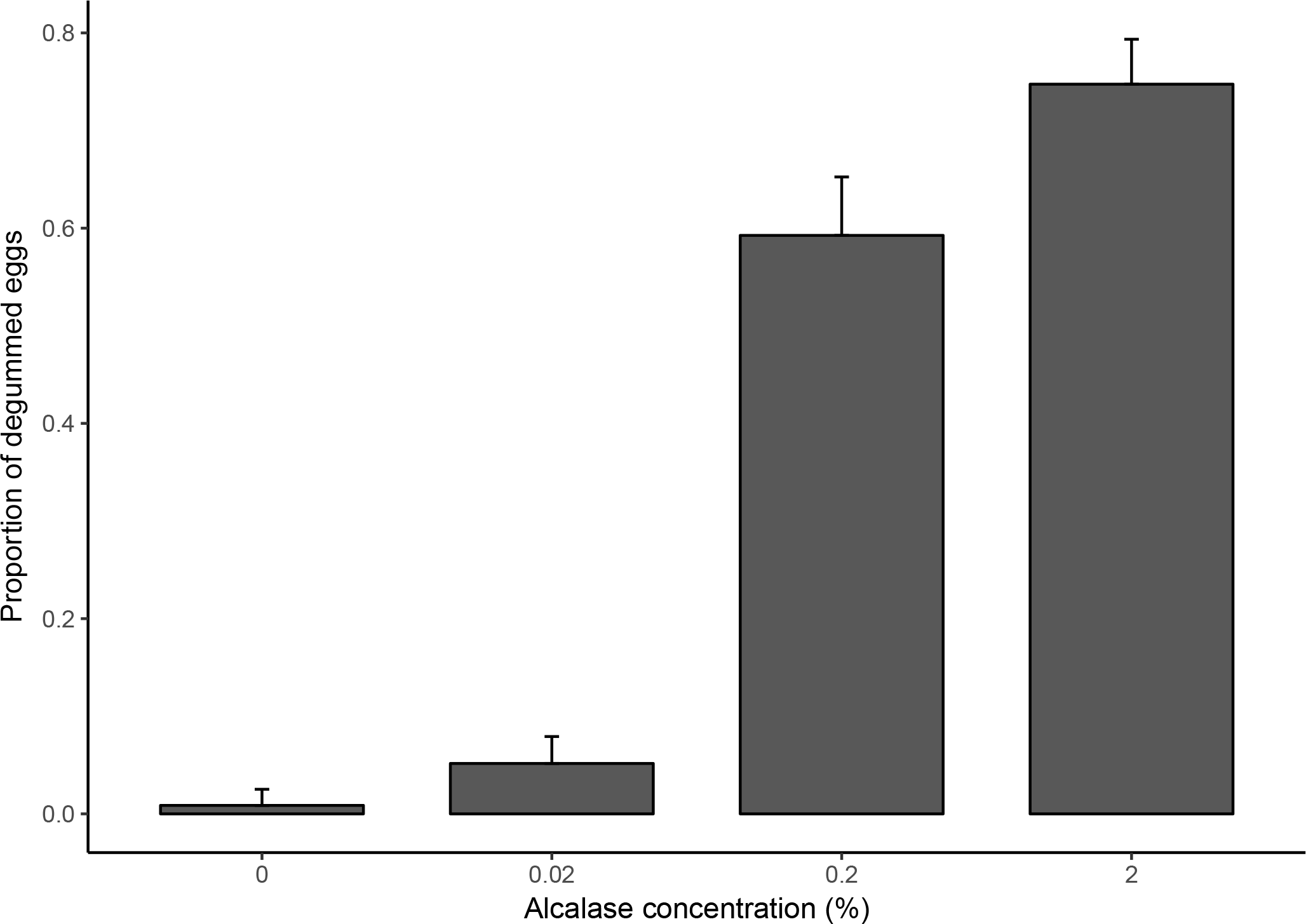
Effect of a 5-minute exposure to different alcalase concentrations on proportion (mean ± 95% CI) of degummed lumpfish eggs

**Figure 2.**
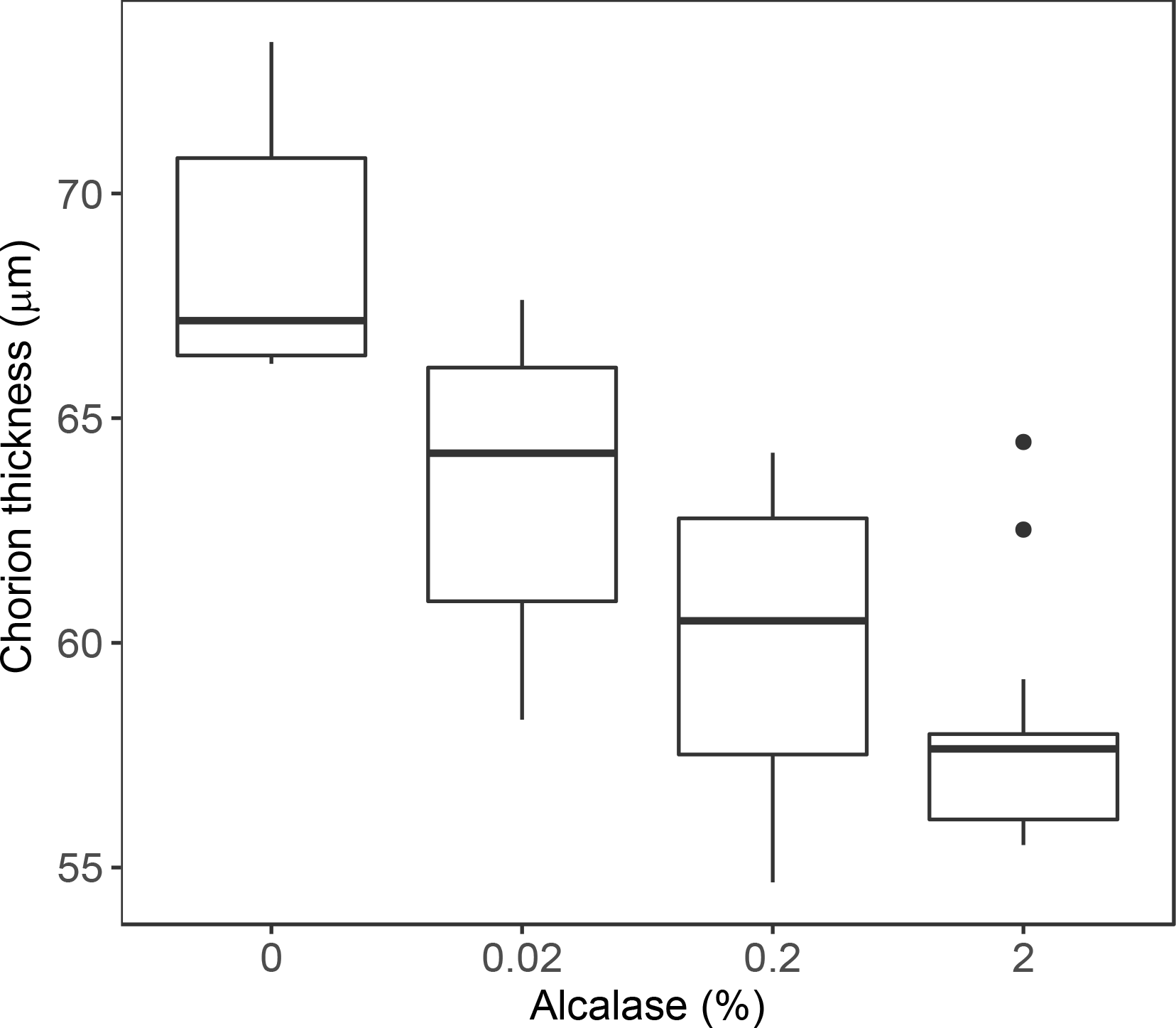
Effect of alcalase treatment on thickness of the egg chorion. Boxplots show minimum, first quartile, median, third quartile, and maximum values. Extreme values are shown by closed circles.

### 3.2. Effect of alcalase on hatching rate and timing of hatching

There was no significant difference in hatching success between controls and eggs exposed to alcalase, once tank effects were taken into account (Figure 3; frailty model treatment estimate = −0.084, SE = 0.056, Z = −1.48, *P* = 0.138; Exponential distribution Loglik(model)= −2801.9, Loglik(intercept only)= −2806.4; χ^2^ = 8.95, df = 4.2, *P* = 0.07). However, alcalase treatment resulted in a shorter hatching period compared to controls (*F*_3,754_ = 69.55, *P* < 0.001), decreasing mean hatching time by −0.66 days (± 0.11) in the case of 0.02% alcalase (*t* = −5.974, *P* < 0.001) and by −1.59 days (± 0.11) in the case of 2% alcalase (*t* = −14.097, *P* < 0.001).

**Figure 3.**
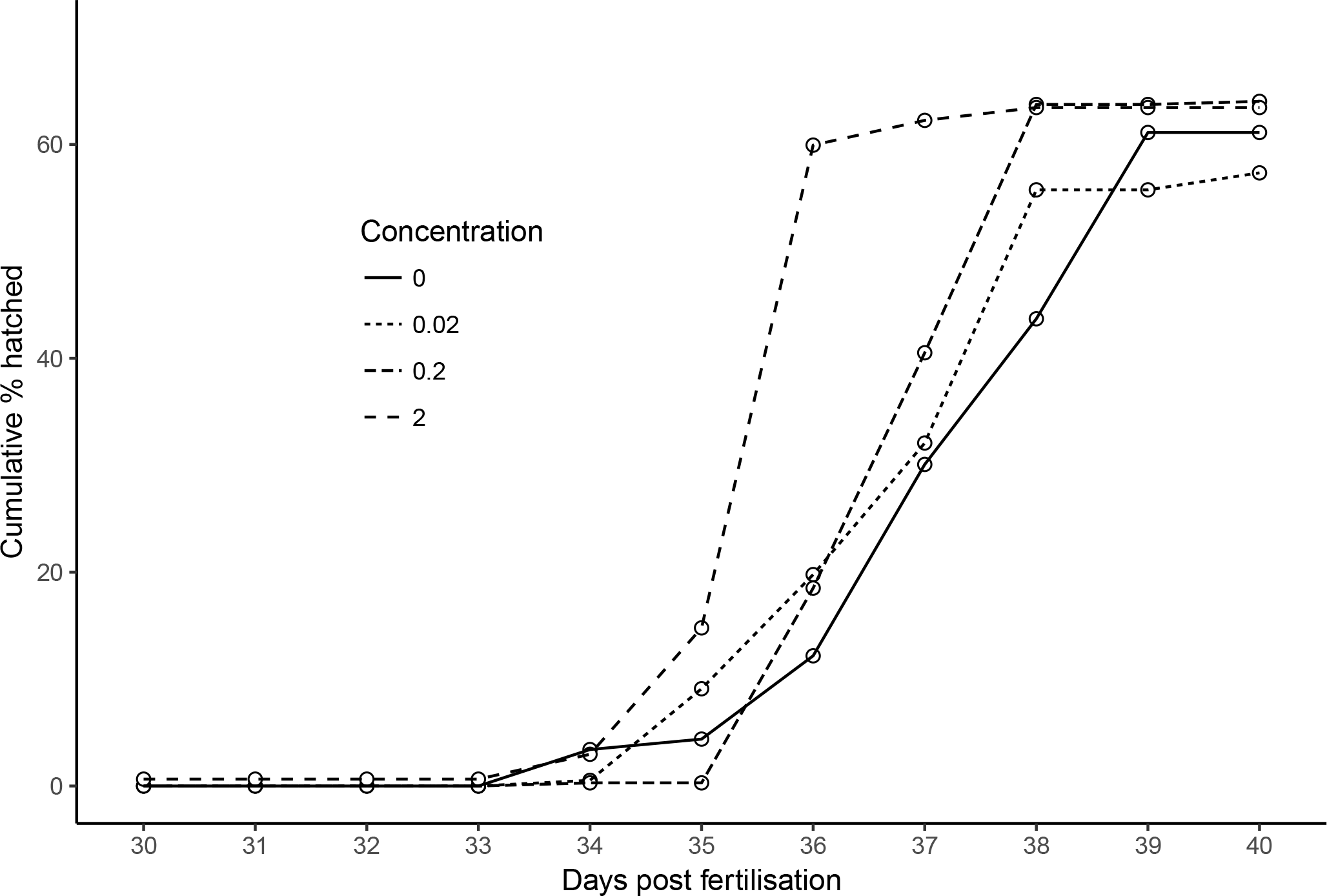
Effect of alcalase treatment on hatching success and timing of hatching (cumulative proportion of hatched larvae).

### 3.3. Effect of alcalase on larvae size and incidence of malformations

Embryo size increased with increasing alcalase concentration (Figure 4, lmer: estimate = 53.99, SE = 19.27, *t*_69.03_ = 2.80, *P* = 0.006). Compared to controls, eggs treated with 2% alcalase resulted in embryos that were 148.14 ± 44.76 µm larger (*t*_68.98_ = 3.31, *P* = 0.001). No deformities were observed under the microscope in any of the larvae from any treatment.

**Figure 4.**
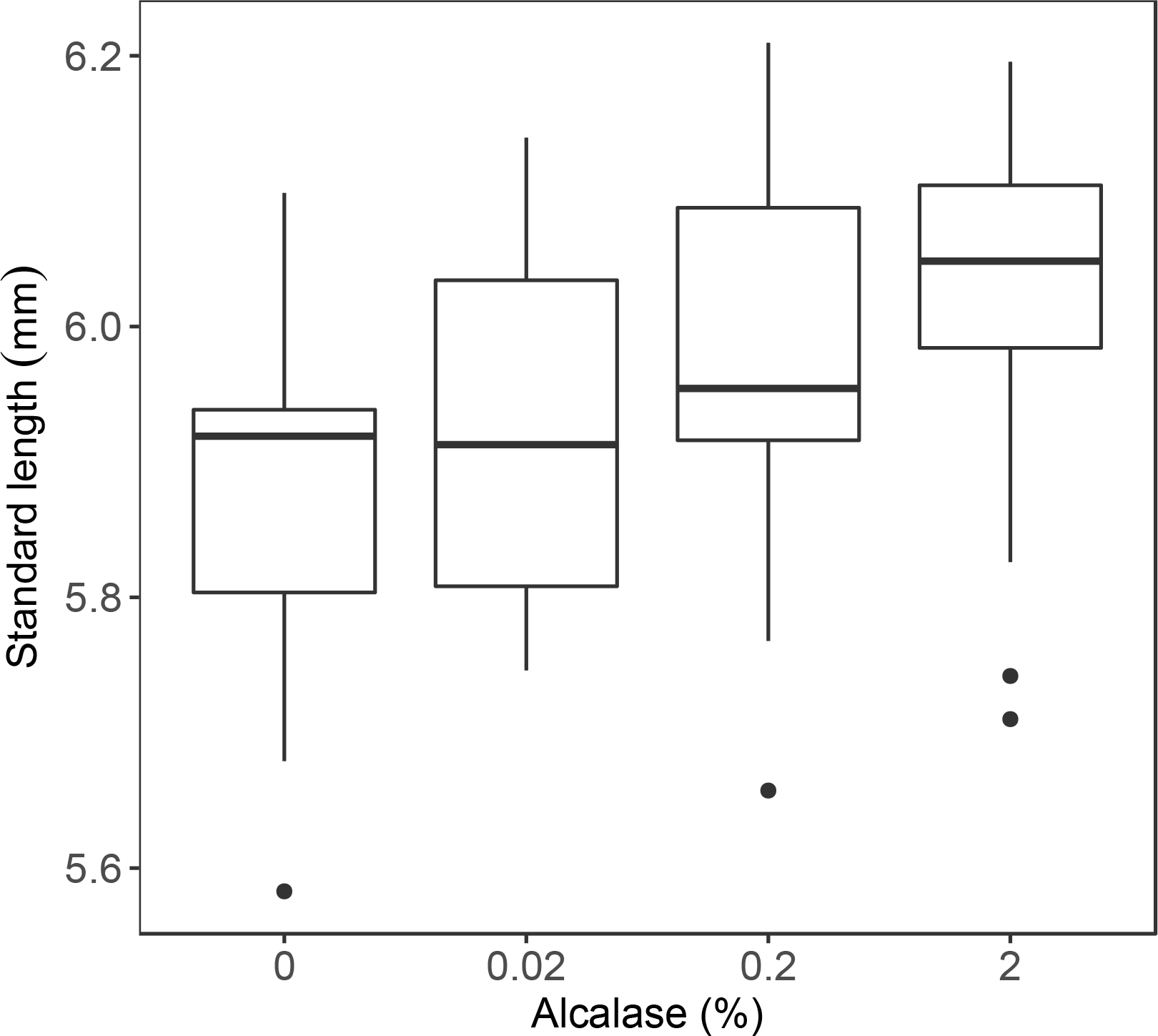
Effect of alcalase treatment on size of lumpfish larvae (standard length). Shown are minimum, first quartile, median, third quartile, and maximum values. Extreme values are shown by closed circles.

### 3.4. Effects of alcalase under commercial conditions

Alcalase was also effective at degumming lumpfish eggs under large scale, commercial conditions (estimate = 0.531, SE = 0.230, Z value = 2.306, *P* < 0.05). Exposure of fertilised eggs to 2% alcalase for five minutes resulted in a mean degumming rate of 0.70 ± 0.06, compared to 0.0 for controls. Alcalase treatment had no significant effect on larvae survival eight weeks post-hatch (estimate = −0.2285, SE = 0.1782, *t* = −1.282, *P* = 0.2), but eggs exposed to 2% alcalase were significantly larger (mean = 14.4 mm ± 0.14) than controls (mean = 13.3 mm ± 0.15; *F*_1,239_ = 41.62, *P* < 0.001), confirming the results of the *in vitro* laboratory trial.

### 3.5. Cost benefit analysis

During the commercial trial, we exposed fertilised lumpfish eggs at a ratio of one egg to one millilitre of alcalase solution. The alcalase solution was purchased at a cost of £46.45 for 500ml, but placing a large bulk order can probably obtain it c. 25% cheaper at £74.3 per Litre. Based on this bulk cost, and £10,000 fixed costs for the sourcing of wild broodstock, we estimated the benefit to cost ratio of incubating 1 million lumpfish eggs under different egg market prices (1-8 pence/egg), taking into account different hatching and degumming rates at different alcalase concentrations. The simulations (Figure 5) suggest that the optimal alcalase concentration is 0.2% (over five minutes), and that benefits outweigh costs at 1.56 pence per egg which is the break even point.

**Figure 5.**
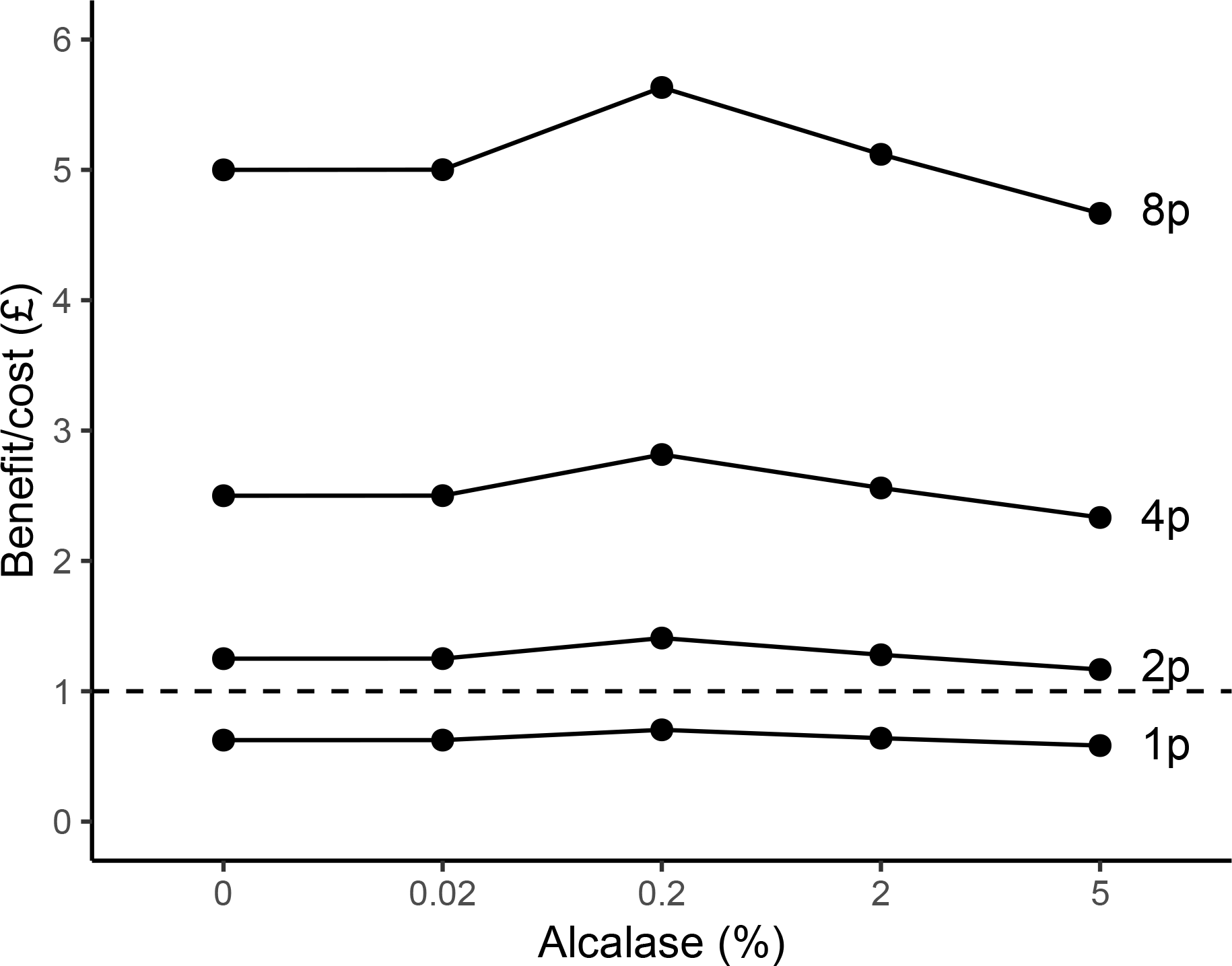
Benefit-cost analysis for the production of 1 million lumpfish eggs using different alcalase concentrations and various egg sale prices (GBP pence/egg). Dotted line represents the estimated break-even point (1.56 pence/egg).

## 4. Discussion

Without parental care in hatcheries, adhesive eggs tend to display reduced gaseous exchange rates, and this can make them more susceptible to opportunistic pathogens (Demska-Zakes et al., 2005). Our results show that alcalase can be effective at removing the adhesive layer of the lumpfish chorion, and that treated eggs result in larger larvae, without any evidence of adverse effects on larval survival or embryo malformations. These findings are similar to work carried out on ballan wrasse, where alcalase treatment achieved >69% degumming rate on a dosage-dependent fashion without any detrimental effects (Grant et al., 2016).

The advantages of egg degumming are various. Degummed eggs allow for more thorough disinfection, easier handling, more accurate egg counting and estimation of fecundity and survival rates, and potentially also more efficient use of space for egg incubation. This can in turn increase hatching success, reduce health risks, and allow for more efficient commercial production (Linhart et al., 2000; Linhart et al., 2003).

Our results indicate that although a five-minute exposure to 2% alcalase achieved the highest degumming rate, a concentration of 0.2% achieved similar results and can be more cost effective, based on a cost-benefit analysis. Little degumming, on the other hand, was observed at a concentration of 0.02% alcalase, although it is possible that increasing exposure time over 5 minutes may achieve better degumming rates, as shown for ballan wrasse (Grant et al., 2016). We exposed eggs to alcalase within ten minutes of artificial fertilisation, before the eggs had hardened. Further research into the feasibility of degumming naturally spawned (hardened) eggs may be warranted, as naturally spawned clumps of eggs are often encountered in culture (Treasurer et al., 2018).

One unexpected consequence of alcalase treatment was a faster embryo development time and larger larvae. Although the reasons for this are unclear, it is possible that the thinner chorion of degummed eggs may have increased oxygen availability and allowed for more efficient use of energy. Oxygen availability is known to affect egg quality and embryo development in carp (Vodianitskiy et al., 2017), and in general increased oxygen results in larger embryos for many fish species (Braga Goncalves et al., 2015). Embryos developing within thick gelatinous egg masses often show retarded development in marine gastropods (Chaffee & Strathmann, 1984), and it is possible that the same may happen in lumpfish. Whatever the reasons, a significant size advantage was evident 8 weeks post-hatching, even though degummed eggs only hatched less than 2 days earlier than controls. The larger size of larvae originating from degummed eggs could be beneficial to industry, as larger larvae have a greater probability of surviving past the critical period during the transition from endogenous to exogeneous feeding (Garrido et al., 2015).

Although a thick egg chorion can make it more difficult for water moulds, such as *Saprolegnia,* to penetrate the eggs (Songe et al., 2016), it can also favour settlement of water borne particulates that facilitate fungal and bacterial infection and can make eggs more vulnerable. Indeed, an adhesive chorion is thought to be particularly susceptible to viral, fungal and bacterial infection, and one of the benefits of achieving a thinner chorion through chemical degumming may be to reduce pathogen loads and facilitate egg prophylaxis under aquaculture conditions (Krise et al., 1986). Egg degumming in lumpfish could also make it possible to use various types of upwelling incubators that maintain eggs in suspension, such as McDonald-type jars, pelagic egg jars, and Imhoff cones (i.e. Jensen et al., 2008). Their use could achieve a more uniform distribution of clean, oxygenated water with less chance of infection by water moulds. This is an area that warrants further attention because growth of opportunistic bacteria and water moulds on dead eggs can spread quickly and affect surrounding viable eggs (Davenport, 1983). Removing the adhesive layer can help to alleviate this problem as it would enable easier separation and removal of non-viable eggs. In this sense, several techniques are available for removing and counting eggs, including manual siphoning, use of flotation methods (i.e. Leitritz & Lewis, 1976) and use of automatic egg sorters, which could prove effective at separating large numbers of eggs if these are degummed and no longer form clumps. For example, using the method described by Coombs (1981), viable and non-viable lumpfish eggs could be separated using a column and a salinity gradient.

## Conclusions

In summary, we describe (and validate under commercial conditions), a method for chemically removing the adhesive layer of lumpfish eggs immediately after artificial fertilization using the enzyme alcalase. A five minute exposure to 2% alcalase resulted in 70% degumming rate, 14% reduction in chorion thickness, and larger larvae without impacting on hatch rates or larvae survival up to 8 weeks after hatching. Cost-benefit analysis indicates that the costs of alcalase treatment are offset under most conditions and that a 0.2% concentration is the most cost-effective. Egg degumming can facilitate the separation and removal of non-viable eggs, improves egg husbandry, and provides scope for using modified incubators that can reduce incubation space and decrease labour time compared to current methods.

## Acknowledgments

This research was part-funded by Marine Harvest Scotland and the ERDF SMARTAQUA Operation. The funders played no role in the collection, analysis or interpretation of the data, the writing of the MS, or the decision to submit the article for publication. We are grateful to staff at CSAR for helping with the experimental set up and egg husbandry, and to Adam Powell and Maria Scolamacchia for setting an initial pilot trial.

